# Analyzing Performance of Twist Bioscience Exome Enrichment with Spike-in CNV Backbone Panels at Various Probe Densities Leveraging Golden Helix VS-CNV Analysis Software

**DOI:** 10.1101/2024.05.19.594885

**Authors:** Nathan Fortier, Gabe Rudy, Tina Han, Alessandro Davassi, Andreas Scherer

**Affiliations:** Golden Helix Inc, 1487 14^th^Ave, Bozeman, MT 59715, USA; Twist Bioscience, 681 Gateway Blvd, South San Francisco, CA 94080, USA

## Abstract

Clinical Whole Exome Sequencing (WES) offers a high diagnostic yield test by detecting pathogenic variants in all coding genes of the human genome. WES is poised to consolidate multiple genetic tests by accurately identifying Copy Number Variation (CNV) events, typically necessitating microarray analysis. However, standard commercial exome kits are typically limited to targeting exon coding regions, leaving significant gaps in coverage between genes, which could hinder comprehensive CNV detection. To convert microarray CNV calling with NGS, advances in both assay design and computational methods are needed.

Addressing the need for comprehensive coverage, Twist Bioscience has developed an enhanced Exome 2.0 Plus Comprehensive Exome Spike-in panel with added CNV “backbone” probes. These probes target common SNPs polymorphic in multiple populations and are evenly distributed in the intergenic and intronic regions, with three varying densities at 25 kb, 50 kb, and 100 kb intervals from highest to lowest resolution respectively. Concurrently, Golden Helix has developed a multi-modal CNV caller designed specifically for target-capture NGS data to detect single-exon to whole-chromosome aneuploidy CNV events. This study evaluates the combined efficacy of the backbone-probe enhanced exome capture kit and VS-CNV 2.6 in identifying known CNVs using the Coriell CNVPANEL01 reference set.

The integration of the enhanced capture kit with VS-CNV 2.6 achieved a 100% sensitivity rate for the detection of known CNV events at all three probe densities. The application of best-practice quality metrics, annotations, and filters was shown to have a minimal impact on this high sensitivity. These findings underscore the potential of the augmented Twist Exome in tandem with the VS-CNV caller and VarSeq’s annotation and filtering capabilities. This combination presents a promising alternative to conventional microarray assays, potentially consolidating WES and CNV into a single assay obviating the need for additional testing in clinical CNV detection. The study’s results advocate for the implementation of this integrated approach as a more efficient and equally sensitive method for CNV analysis in a clinical setting.

## Introduction

Copy Number Variants (CNVs) are a common source of genomic variation and have been implicated as causal mutations in a wide variety of human diseases [1]. The first generation of genetic tests for CNV detection relied on cytogenetic technologies and included karyotyping through direct inspection of the count, length, and shape of chromosomes, as well as fluorescence in situ hybridization (FISH) [2]. In later years, chromosomal microarray analysis (CMA) emerged as a powerful tool for the detection of large CNVs [3]. However, CMA technology is unable to detect small intragenic CNVs that only span a portion of a gene. These CNVs can have a significant clinical impact but are necessarily missed by even the highest density microarrays. To address this limitation, many clinical labs have adopted multiplex ligation-dependent probe amplification (MLPA) as the gold standard for the detection of smaller intragenic CNVs [4]. While MLPA excels at this use case, the cost of this technology increases with the number of genes targeted, and it is unable to detect the large cytogenetic events that would typically be detected using CMA technology.

To address the limitations of CMA and MLPA technologies, Golden Helix has developed the VS-CNV algorithm as part of the VarSeq Software Suite [5]. This algorithm detects CNVs based on Next-Generation sequencing (NGS) data and is capable of detecting CNVs ranging in size from small intergenic events to large multi-gene CNVs. Because NGS technology is already widely used for the detection of single nucleotide variations (SNVs), VS-CNV has the added benefit of enabling the detection of both CNVs and SNVs using a single assay.

However, CNV detection is only the first step of the CNV analysis workflow. Performing CNV calling on a single whole exome or whole genome sample will typically result in hundreds of CNV calls, making the task of manually interpreting each event practically infeasible. Thus, tertiary analysis workflows must be developed to filter out false positive CNV calls and identify clinically relevant CNVs. This requires a combination of using CNV-specific QC values to remove low-quality CNV calls and annotating CNV calls with gene impact analysis and relevant clinical annotations.

To support this need, Golden Helix has developed a robust set of tools for annotating and filtering CNVs in the VarSeq Software Suite. One of these tools is an algorithm that automates the application of many criteria specified in the 2020 Guidelines for CNV scoring published by the ACMG and ClinGen [6]. These guidelines extend the 2015 ACMG Guidelines for the interpretation of sequence variants published by Richards *et al*. [7] to provide guidance on the interpretation of CNVs. This algorithm is integrated into VarSeq and can be used to filter out benign and likely benign CNVs, while prioritizing CNVs with evidence of pathogenicity. It provides a classification of each CNV based on the classification tiers specified in the ACMG Guidelines, along with gene impact scores that quantify the predicted impact of the CNV on the gene product [8].

While VS-CNV is commonly used in conjunction with standard commercial exome kits, whole exome sequencing technology is limited to targeting exon coding regions, leaving significant gaps in coverage between genes, which could hinder comprehensive CNV detection. To address the need for comprehensive coverage across intergenic regions, Twist Bioscience has developed an enhanced Exome 2.0 Plus Comprehensive Exome spike-in panel with added “backbone” probes. These probes are evenly distributed in the large intronic and intergenic regions, with three varying densities at 25 kb, 50 kb, and 100 kb intervals.

In this paper, we evaluate the performance of VarSeq’s CNV calling and filtering capabilities in conjunction with the backbone-probe enhanced exome capture technology developed by Twist Bioscience. Our experiments quantify the sensitivity of both the VS-CNV algorithm and a best-practice tertiary filtering workflow in VarSeq, which incorporates quality metrics along with classification and scoring information provided by the ACMG CNV Classifier to identify high-quality CNVs with evidence of pathogenicity. This evaluation was performed on 42 samples with known CNVs and disease states. These benchmark samples were sequenced using the Twist Exome 2.0 Plus Comprehensive Exome Spike-in and the Twist CNV Backbone Spike-in Panel, which can be combined to capture both the WES and CNV targets within a single enrichment capture. The sensitivity of the VS-CNV algorithm and the proposed filtering workflow were evaluated using a benchmark dataset of confirmed CNVs provided by the Coriell Institute for these samples. Each sample was sequenced at three different levels of probe density to assess how changes in probe density affect the performance of the CNV analysis workflow.

## Experimental Design

### Review of the Study Data

To evaluate the performance of VS-CNV and the associated filtering workflow, we have called CNVs from NGS data for 42 of the samples in the Copy Number Variation Panel from the Coriell Institute’s NIGMS Human Genetic Cell Repository [9]. Each sample in the panel harbors at least one clinically significant chromosomal aberration. The Coriell Institute provides a manifest of the confirmed CNV and loss of heterozygosity (LoH) events associated with each sample as well as the diagnosed disorder from the clinical evaluation or publication that sourced the sample. While this dataset does not provide a comprehensive list of all detectable CNVs that may be present in each sample, all events in this dataset have been confirmed through G-banded karyotyping analysis and, in many cases, FISH. This benchmark dataset is the basis for establishing the sensitivity of the VarSeq CNV calling and filtering workflow. The set of 55 CNVs used in this analysis, along with their associated samples and disorders are shown in Table 1. These events range in size from 100,843 bp to 155,000,000 bp (100 kb to 155 mb).

**Table 1:** Coriell Institute Benchmark CNVs.

| Sample ID | Disorder | CNVs |
| --- | --- | --- |
| GM01416 | XXXX Syndrome | X:251797-155952677 Gain |
| GM05067 | Aneuploid Chromosome Number - Trisomy 9 | 9:46586-39778619 Gain |
| GM05966 | Derivative Chromosome | 14:54502047-75681317 Gain |
| GM06226 | Translocated Chromosome | 1:247143229-248930177 Loss<br>16:35814-21378649 Gain |
| GM06870 | Aneuploid Chromosome Number - Non-Trisomic | 18:11542-15401752 Gain |
| GM06936 | Chromosome Deletion | 10:58486-12836926 Loss |
| GM07945 | Adenosine Deaminase Deficiency With No Immunodeficiency | 13:20228923-20460156 Loss<br>20:34910451-46231832 Loss |
| GM08331 | Chromosome Deletion | 13:97506715-109611222 Loss<br>21:25943808-28146869 Loss |
| GM09102 | Chromosome Deletion | 11:120620269-135074876 Loss |
| GM09216 | Chromosome Deletion | 2:10203410-26929010 Loss<br>4:143920938-144021781 Loss |
| GM09367 | Duplicated Chromosome | 6:107433158-142743017 Gain |
| GM09888 | Trichorhinophalangeal Syndrome, Type II | 8:106107809-118282364 Loss<br>14:50355382-51188531 Loss |
| GM10608 | Chromosome Deletion | 20:9891954-18050825 Loss |
| GM10636 | Duplicated Chromosome | 2:109704906-110869845 Gain<br>X:39949336-57930356 Gain |
| GM10800 | Chromosome Deletion | 4:69196130-94156942 Loss |
| GM10925 | Greig Cephalopolysyndactyly Syndrome | 7:38592416-54646811 Loss<br>X:372539-654184 Gain |
| GM10985 | Chromosome Deletion | 3:18654-10288693 Loss |
| GM10989 | Gilles De La Tourette Syndrome | 9:46586-11996831 Loss |
| GM11213 | Chromosome Deletion | 2:186245475-203738206 Loss |
| GM11419 | Aneuploid Chromosome Number - Non-Trisomic | 4:143899562-144129458 Gain<br>Y:2782384-26653776 Gain |
| GM11672 | Chromosome Deletion | 10:48084407-73690949 Loss |
| GM12606 | Chromosome Deletion | 13:18471487-59667287 Gain |
| GM12662 | Chromosome Deletion | Y:22213783-26541710 Loss |
| GM13019 | Turner Syndrome | X:251797-56869151 Loss<br>X:63123908-154822179 Gain |
| GM13464 | Williams-Beuren Syndrome | 7:73311765-74727754 Loss |
| GM13476 | Smith-Magenis Syndrome | 17:16860240-20492562 Loss |
| GM14164 | Tetralogy Of Fallot | 13:47227948-95062722 Loss |
| GM14485 | Inverted Duplication Deletion | 8:220289-7368769 Loss<br>8:12670906-43745225 Gain |
| GM14943 | Chromosome Deletion | 2:234368396-242147293 Loss |
| GM16362 | Aneuploid Chromosome Number - Trisomy | 22:16986520-19090414 Gain<br>22:21960704-22219583 Loss<br>22:42883607-50796015 Gain |
| GM16595 | Cri-du-cha Syndrome | 5:8633692-24036533 Loss |
| GM17867 | Klinefelter Syndrome | X:251797-155699825 |
| GM17942 | DiGeorge Syndrome | 22:18167914-21108322 Loss |
| GM20022 | Duplicated Chromosome | 3:134843252-195926102 Gain |
| GM20027 | Turner Syndrome | X:251797-156004181 Loss |
| GM20556 | Isodicentric Chromosome | 15:19811062-32478247 Gain |
| GM21698 | Chromosome Deletion | 6:162519205-170610395 Loss |
| GM21699 | Chromosome Deletion | 3:18654-564371 Gain<br>6:163241020-170673434 Loss |
| GM21887 | Angelman Syndrome | 15:17167687-23199681 Loss |
| GM22601 | Wolf-Hirschhorn Syndrome | 4:65772-25980331 Loss |
| GM22624 | Potocki-Shaffer Syndrome | 11:40455217-46053197 Loss |
| GM22991 | Chromosome 1P36 Deletion | 1:817185-5250920 Loss |

An additional 13 samples from the Coriell Institute that are not known to harbor any large CNVs were also included as supplementary reference samples. All 55 samples were included in the reference set for CNV calling. Because there is little overlap between the locations of the various CNVs across the CNV panel samples, including the 42 non-normal benchmark samples as references is unlikely to skew the results.

### CNV Workflow Evaluation Design

One goal of this study is to confirm that these CNVs can be accurately detected using NGS technologies. To evaluate the sensitivity of CNV detection, the Coriell Institute samples have undergone the following NGS primary and secondary analysis steps:

1. Fifty nanograms of gDNA were subjected to the Twist Library Preparation Enzymatic Fragmentation 2.0 Kit. Unique dual indices (UDIs) were added during pre-capture PCR. Multiple sample libraries were pooled together for an overnight hybridization with Twist Exome 2.0 Plus Comprehensive Exome Spike-in and Twist CNV Backbone Spike-in Panels with 25 kb, 50 kb, or 100 kb spacing.
2. Samples were pooled by equal molar ratio and sequenced on Illumina NovaSeq 6000 as paired-end 150 reads (PE150). Raw sequencing data as fastq file were demultiplexed using UDIs with Illumina bcl2fastq (2.20) and 1 mismatch was allowed.
3. Alignment and Variant Calling was performed using the Sention DNAScope pipeline version 202308.
4. The VS-CNV algorithm was run on each batch of samples of the same probe density using the default algorithm settings in VarSeq 2.6.0.

### CNV Count Statistics

We report the total number of CNVs called along with the number of filtered CNVs at each step of the tertiary analysis workflow. These CNVs are divided into two size categories:

1. CNVs larger than 10 kb
2. CNVs smaller than 10 kb

The CNVs falling into the first size category are consistent with the size of events that are detectable using a typical microarray platform, such as the Agilent CGH + SNP array [10]. CNVs falling into the second category would not be detectable via traditional microarray technology. For each size category, the results of this study report the following statistics:

- Total number of called CNVs.
- Number of high-quality CNVs.
- Number of CNVs with evidence of pathogenicity.

### High Quality CNVs

The quality of each CNV was assessed using the calculated p-value and the assigned quality flags. The p-value estimates the probability that z-scores at least as extreme as those in the event would occur by chance in a normal region and is computed automatically for each CNV using a paired Student’s t-test. For these results, we consider a CNV call to be high quality if it has a p-value < 0.01 and has not been assigned any quality flags other than Extreme GC Content. We have excluded this flag from the filter chain, as some of the CNVs in the benchmark dataset are in regions that are known to have extreme GC content. This flag is intended to notify users when CNVs are in such regions, as these regions typically have poor mappability resulting in less reliable CNV calls, but the flag does not necessarily indicate a low-quality call.

### CNVs with Evidence of Pathogenicity

A CNV is considered to have evidence of pathogenicity if the associated output from the ACMG CNV Classifier meets the following criteria:

- The CNV is not classified as Benign / Likely Benign.
- The CNV is classified as Pathogenic / Likely Pathogenic or has a *Potential Gene Impact Score* that is either missing or greater than or equal to 0.9.

The *Potential Gene Impact Score* provides the score a CNV would receive in accordance with the ACMG Guidelines if its overlapping genes were known to be haploinsufficient or triplosensitive. The threshold of 0.9 indicates that the CNV would be classified as Pathogenic / Likely Pathogenic if the gene of interest was shown to be haploinsufficient or triplosensitive. For large CNVs overlapping more than 50 genes, this score is not computed, so CNVs that have a missing value for this score should not be excluded by the filtering workflow.

### Sensitivity of the CNV Caller

Due to the granular nature of CNV calling on NGS coverage data, large cytogenetic events may be called as several distinct smaller events overlapping the region. Additionally, the targets used to call the CNV may not perfectly overlap with the region defined in the Coriell Institute’s benchmark dataset. Based on these considerations, each analyzed CNV in the benchmark dataset will be categorized as either a True Positive or False Negative as follows:

- True Positive: More than 75% of targets in the benchmark region have been assigned the correct state by the CNV Caller.
- False Negative: Fewer than 75% of targets in the benchmark region have been assigned the correct state by the CNV Caller.

The number of CNVs falling into these categories will then be used to compute the sensitivity of the VS-CNV algorithm as follows:

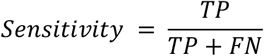

### Sensitivity of the Filtering Workflow

To evaluate the sensitivity of each step of the filtering workflow, each CNV in the benchmark dataset will be categorized as either a True Positive or False Negative as follows:

- True Positive: The benchmark CNV overlaps a called CNV in the filtered results with the appropriate CNV State.
- False Negative: The benchmark CNV does not overlap any called CNV in the filtered results with a matching CNV State.

The number of CNVs falling into these categories will then be used to compute the sensitivity of each step of the filtering workflow:

1. **Quality Filtering Step**: Filter to high-quality CNVs that have a p-value < 0.01 and have not been assigned any quality flags other than Extreme GC Content.
2. **Classifier Filtering Step**: Filter to CNVs with evidence of pathogenicity as indicated by the automatic ACMG Classification and Potential Gene Impact Score.

This filtering workflow is designed to include only high-quality CNVs with evidence of pathogenicity as defined in the discussion of CNV Count Statistics. A complete description of the specific filters used in this workflow can be found in the appendix.

### Precision of the Filtering Workflow

Because the benchmark dataset provided by the Coriell Institute is not comprehensive, it is not possible to directly measure the precision as there is no way to discern false positive CNV calls from true events that are simply not present in the benchmark data. However, the total number of CNVs falling into the greater than 10 kb size category at each step of the filtering process provides a proxy measure of the workflow’s precision. The sensitivity of the VS-CNV algorithm is 100% across all probe densities, with all events in the benchmark dataset being called over at least 75% of overlapping targets. The quality filtering workflow does not have any impact on sensitivity for the 100 kb probe density, with no true positive events being filtered out at this probe density. However, the 25 kb and 50 kb probe densities result in a slightly reduced sensitivity for the quality filtering workflow, with two benchmark CNVs filtered out due to being flagged as having *High Controls Variation*. This flag is applied when the coefficient of variation for the normalized coverage across the called targets exceeds 0.25 and indicates a high degree of variation between the reference samples for the region of interest. The two filtered benchmark CNVs are described below:

- Sample GM10925 - X:372539-654184 Gain
- Sample GM12662 - Y:19928754-26212843 Loss

### Probe Density

A secondary goal of this study is to evaluate the impact of probe density on performance given the above study design. Probe density refers to the genomic distance between the probes used to target and capture genomic regions of interest. The end-point precision of CNV calls will depend on the density of the probes, with higher probe density allowing for more precise determination of CNV boundaries. The samples in this panel have been sequenced at three levels of probe density:

1. 25 kb
2. 50 kb
3. 100 kb

We have evaluated the performance of the VarSeq CNV workflow using the above study design at each probe density level to assess how changes in probe density affect the performance of the CNV analysis workflow.

## Experimental Results

### Sensitivity of CNV Caller and Filtering Workflow

The sensitivity of both the VS-CNV algorithm and the filtering workflow for each level of probe density is shown in Table 2.

**Table 2:** Sensitivity of Caller and Filtering Workflow.

| Probe Density | 25 kb | 50 kb | 100 kb |
| --- | --- | --- | --- |
| VS-CNV Sensitivity | 100% (55/55) | 100% (55/55) | 100% (55/55) |
| Quality Filtering Sensitivity | 96.4% (53/55) | 96.4% (53/55) | 100% (55/55) |
| Classifier Filtering Sensitivity | 94.5 (52/55) | 94.5% (52/55) | 98.2% (54/55) |

The normalized coverage of the targets in these regions appears to have much more variation across samples for the higher probe densities, which results in these CNVs being flagged. This may indicate that the backbone probe regions exhibit higher variation in coverage between samples resulting in increased coverage variably at higher probe densities. VarSeq offers tools for identifying such highly variable regions and it is possible that excluding these regions from the analysis could result in improved sensitivity. For future work we will evaluate the viability of such target filtering strategies.

The classifier filtering workflow does result in slightly decreased sensitivity (98.2% for the 100 kb probe density), with a single additional event being filtered out at all probe densities. This event is a deletion of the last four exons of the gene GYPB in sample GM09216, spanning the region 4:143920938-144021781. This CNV was classified as Benign due to the application of criterion 4O of the ACMG Guidelines, as the gene’s coding region is fully contained by a high frequency GnomAD [11] CNV region with an average frequency of 0.04 and a high frequency DGV [12] CNV with an allele frequency of 0.01. GYPB is a red-cell protein but one that is not essential for red-cell development or human survival [13]. Therefore, loss of this gene is tolerated and the CNV filtering workflow **is correct** in removing this benign CNV.

Based on these results, a conclusion can be made on the question of probe density. For the samples in this experiment, probe density **does not impact** the sensitivity of the VS-CNV algorithm. However, the higher probe densities of 25 kb and 50 kb do result in reduced sensitivity for the quality filtering workflow due to increased variation in normalized coverage between reference samples. This result is unsurprising as all events in the benchmark dataset were larger than 100 kb in size. It is possible that the algorithm’s sensitivity for the detection of smaller CNVs would be impacted by increased probe density.

### CNV Count Statistics

The CNV count statistics for each size category and probe density are shown in Table 3. These counts include all CNVs across the 42 samples with confirmed CNVs in the benchmark dataset.

**Table 3:** CNV Count Statistics for Benchmark Samples.

| Probe Density | 25 kb |  | 50 kb |  | 100 kb |  |
| --- | --- | --- | --- | --- | --- | --- |
| Event Size | > 10 kb | < 10 kb | > 10 kb | < 10 kb | > 10 kb | < 10 kb |
| Called CNVs | 8,787 | 23,247 | 5,759 | 17,756 | 4,825 | 16,338 |
| High-Quality CNVs | 2,672 | 2,397 | 1,683 | 1,854 | 1,398 | 1,683 |
| CNVs with Evidence of Pathogenicity | 706 | 621 | 482 | 475 | 434 | 473 |

Both the total number of CNVs called by the algorithm and the number of high-quality filtered CNVs increase as the distance between the probes decreases. The majority of the called events across all probe densities are below the 10 kb size threshold, but most of these events are either low-quality or do not have evidence of pathogenicity. This is unsurprising as the events spanning more than 10 kb generally span large numbers of targets, resulting in more reliable calls. Additionally, most of these events fully contain multiple genes and are therefore more likely to have evidence of pathogenicity. The average number of CNVs per sample in each size category is presented in Table 4 below.

**Table 4:** Average Number of CNVs per Sample.

| Probe Density | 25 kb |  | 50 kb |  | 100 kb |  |
| --- | --- | --- | --- | --- | --- | --- |
| Event Size | > 10 kb | < 10 kb | > 10 kb | < 10 kb | > 10 kb | < 10 kb |
| Called CNVs | 209.2 | 553.5 | 137.1 | 422.8 | 114.9 | 389.0 |
| High-Quality CNVs | 63.6 | 57.1 | 40.1 | 44.1 | 33.3 | 40.1 |
| CNVs with Evidence of Pathogenicity | 16.8 | 14.8 | 11.5 | 11.3 | 10.3 | 11.26 |

For the 100 kb probe density, there is an average of around 73 high-quality CNVs per sample. While this is a drastic reduction compared to the hundreds of unfiltered CNV calls per sample, this large number of high-quality filtered CNVs still presents a challenge for manual interpretation, motivating the need for additional filtering to identify clinically relevant events. However, once filtering is performed based on the ACMG Classifier, there are only around 22 clinically relevant CNVs per sample. These results indicate that a tertiary workflow that combines the filtering of high-quality CNVs with prioritization based on the automated application of the ACMG Guidelines will result in a small number of clinically relevant CNVs requiring manual interpretation.

## Conclusion

In this study, we evaluated the performance of Twist Exome 2.0 plus Comprehensive Exome Spike-in paired with a CNV Spike-in capture panel in conjunction with VarSeq’s CNV analysis capabilities. This includes an evaluation of the CNV detection capabilities of VS-CNV when combined with a best-practice annotation and filtering workflow for the identification of high-quality clinically relevant CNVs. This evaluation was conducted using 42 samples from the Coriell Institute’s Copy Number Variation Panel. The sensitivity of both the CNV caller and filtering workflow was evaluated using a benchmark dataset of confirmed CNVs, with the results indicating excellent performance across various probe densities.

The results demonstrated the ability of the VS-CNV algorithm to consistently detect CNVs of varying sizes, with a consistently high sensitivity of 100% across different probe densities. Additionally, the proposed filtering workflow eliminates large numbers of clinically irrelevant and low-quality CNV calls, with minimal impact on the sensitivity of the workflow. For the 100 kb probe density, filtering based on quality metrics had no impact on sensitivity, while a slight reduction in sensitivity was observed for the 25 kb and 50 kb probe densities. Furthermore, filtering based on the output of the ACMG Classifier drastically reduced the number of clinically relevant CNVs while only removing a single event in the benchmark dataset. This event was filtered due to overlap with high-frequency CNVs in multiple population catalogs, and published literature indicates that this event is likely benign.

This research highlights the potential of the augmented Twist WES capture panel with CNV panels in tandem with VarSeq’s CNV calling, annotation, and filtering capabilities. This workflow has been shown to reliably detect CNV events across the 42 benchmark samples while eliminating the vast majority of low-quality and clinically irrelevant CNV calls. This combination of technologies offers a promising alternative to traditional microarray analysis, potentially eliminating the necessity for additional testing in clinical CNV detection.

## Appendix

### CNV Calling Methodology

The VS-CNV algorithm was run on BAM and VCF files sequenced at each level of probe density using the following methodology in VarSeq 2.6.0:

1. Import VCF files and BAM files for all samples into the VarSeq project.
2. Perform Loss of Heterozygosity (LoH) calling in VarSeq.
3. Run the VarSeq CNV Caller using the default algorithm settings.
4. Apply filter chain to include only Deletion, Het Deletions, and Duplications present in the current sample.
5. Export CNV and Coverage Regions tables.
6. Compute performance metrics from exported tables.

The target regions used by the CNV caller consist of the exon boundaries for the clinically relevant transcripts of all RefSeq genes for which the average target mean depth across all samples exceeds 20x.

### CNV Filtering Workflow

The annotation and filtering workflow applied to the CNVs in VarSeq 2.6.0 was as follows:

1. Import VCF files and BAM files for all samples into the VarSeq project.
2. Import the sample manifest containing the associated phenotypes for each sample.
3. Perform Loss of Heterozygosity (LoH) calling in VarSeq.
4. Run the VarSeq CNV Caller using the default algorithm settings.
5. Annotate CNVs using the ACMG Sample CNV Classifier.
6. Run the PhoRank gene ranking algorithm using the extracted phenotypes.
7. Perform CNV filtering using the following filter chain:
  a. Filter on CNV State to include only Deletions, Het Deletions, and Duplications in the current sample.
  b. Filter out all CNVs flagged with any flag other than
  c. Filter out CNVs with a *p-value* ≥ 0.01.
  d. Filter out CNVs classified as Benign / Likely Benign by the ACMG Classifier.
  e. Filter using the ACMG Classifier to include only CNVs meeting at least one of the following criteria:
    i. *Potential Gene Impact Score* ≥ 0.9.
    ii. *Potential Gene Impact Score* is missing.
    iii. Classification is Pathogenic / Likely Pathogenic.

